# *ninaD* regulates cholesterol homeostasis from the midgut which protects against neurodegeneration

**DOI:** 10.1101/2020.11.27.401059

**Authors:** Stéphanie Soulé, Jean-René Martin

## Abstract

Cholesterol is crucial to maintain normal cellular function. In human, it has also been involved in various neurodegeneration processes, as Niemann-Pick and Alzheimer diseases. Recently, we have identified a small nucleolar RNA (*jouvence*) required in the epithelial cells of the gut (enterocytes), and showed that its overexpression extends lifespan. A transcriptomic analysis has revealed a deregulation of several genes in *jouvence* mutants. Among them, *ninaD* encoding a mammalian homolog to class B Scavenger receptor is importantly upregulated. In *Drosophila, ninaD* is required for the uptake of the dietary carotenoid, used for the formation of rhodopsin. Here, we show that *jouvence*-deleted flies are deficient in cholesterol-ester, as well as old flies present neurodegenerative lesions. Restoring *ninaD* mRNA expression level in enterocytes restores the metabolic cholesterol-ester level, prevents neurodegeneration and extends lifespan, revealing a gut-brain axis. Our studies demonstrates that *ninaD* is a central regulator of cholesterol homeostasis and a longevity-promoting factor.

## INTRODUCTION

Cholesterol is an essential cellular component and its homeostasis is required to maintain normal cellular function. Different forms of cholesterol, as free or ester of cholesterol are dynamically and highly regulated. We have reported the genetic and molecular characterization of a small nucleolar RNA (*jouvence*) (*jou*) required in the enterocytes (Soulé et al., 2020). The loss of *jou* shortens lifespan and leads to a intestinal hyperplasia and defects in metabolic parameters (Biteau et al., 2008; 2010). Inversely, targeted expression of *jou* in enterocytes (overexpression) increases lifespan, and prevents hyperplasia in aged flies gut (Soulé et al., 2020). A RNA-Seq analysis of *jou*-deleted adults gut revealed a deregulation in expression level of several genes. Among them, *ninaD* (*neither inactivation nor afterpotential D*) encoding a mammalian homolog to class B Scavenger receptor-like membrane protein (Kiefer et al., 2002), is upregulated. In *Drosophila, ninaD* is required for the uptake and storage of the dietary carotenoid for the formation of rhodopsin (Kiefer et al., 2002; Gu et al., 2004; Yang et al., 2007; Voolstra et al., 2006). The targeted expression of a UAS-ninaD-RNAi specifically in enterocytes, which knockdown the *ninaD* mRNA, restores the mRNA level and consequently result in lifespan extension (Soulé et al., 2020). Here, we demonstrate that *ninaD* in enterocytes, is essential for cholesterol homeostasis. *jou*-deleted flies accumulate an excess of free cholesterol, but a striking decrease of cholesterol-ester. Moreover, *jou*-deleted old flies present neurodegenerative lesions. Restoring the *ninaD* mRNA level in the gut extends lifespan, restore metabolic cholesterol homeostasis and prevents neurodegeneration, underlying a gut-brain axis.

## RESULTS AND DISCUSSION

### Downregulation of *ninaD* in the gut rescues lifespan in *jou-deleted* flies

We have previously reported that loss of *jou* shortens lifespan (Soulé et al., 2020), a phenotype related to the deregulation of several genes. This phenotype is rescued by specifically restoring the mRNA level of *ninaD* in the enterocytes. To consolidate the role of *ninaD* in regulating lifespan, we used other Gal4 drivers with gut-specific expression pattern to express the UAS-ninaD-RNAi. In *jou*-deleted flies, the enterocytes-specific driver Myo1A-GAL4, used to downregulate *ninaD* mRNA level, leads to lifespan extension (Figure 1A). Similarly, using the CG8997-GS line also expressed in the epithelium of the gut, in *jou*-deleted flies, allowing the expression of the ninaD-RNAi only in adulthood after feeding with RU486, also increases lifespan (Figure 1B). These results confirm that rescuing the *ninaD* mRNA in enterocytes, by knockdown, is sufficient for lifespan extension in *jou-*deleted flies, and even when it is restored only in adulthood.

**Figure 1).**
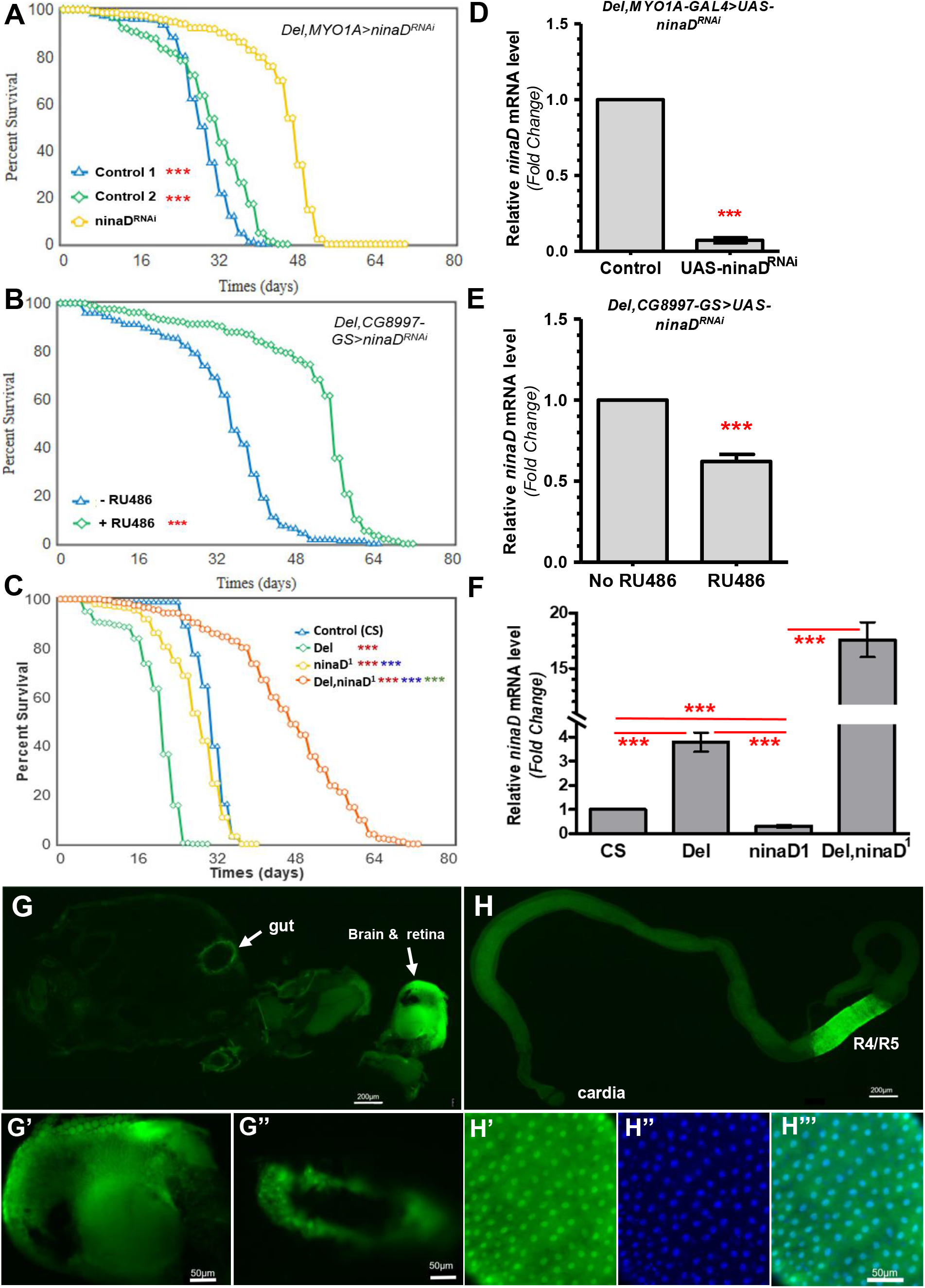
Knockdown of *ninaD* in enterocytes rescues lifespan in *jou-deleted* flies. **(A-C)** Longevity test results (survival curve - decreasing cumulative). **A)** Targeted expression of the UAS-ninaD-RNAi specifically in the enterocytes in *jou*-deleted flies (Del,Myo1A) compared to their two respective controls (Control-1 : Del,Myo1A-Gal4/Del and Control-2 : Del,UAS-ninaD^RNAi^). **B)** Targeted expression, only in adulthood, of the ninaD-RNAi specifically in the enterocytes in Del,CG8997-GS RU486 fed flies compared to their controls (non-fed RU486). **C)** *ninaD^1^* mutant compared to Control (CS), *jou*-deleted (Del), the double-mutant (*Del,ninaD^1^*). Statistics: red=compared to CS, blue=compared to Del, green=compared to *ninaD^1^*. For detailed Statistics, see Suppl. Table S2 (**\*\*\***p<0.001). **D-F)** RT-qPCR of the quantification of the *ninaD* mRNA level. **D)** Del,Myo1A-Gal4>UAS-ninaD-RNAi (correspond to UAS-ninaD-RNAi) compared to their controls (Del,Myo1A-GAL4/Del). **E)** Del,CG8997-GS RU486 fed flies compared to non-fed RU486). **F)** *ninaD^1^* compared to the double-mutant *Del,ninaD^1^*, and to controls (wild-type CS and Del) (***p<0.001). **G-H)** Analysis of expression pattern of *ninaD* using ninaD-Gal4 construct driving expression of the UAS-nls-GFP nuclear marker. **G)** Cryosection of whole fly. The labeling of the retina is nonspecific due to autofluorescence. GFP is expressed in the epithelium of the gut **(G’)** and the brain **(G”)**. **H)** Isolated midgut from ninaD-Gal4>UAS-nls-GFP at 7 days. **H’-H’’’)** Close-up showing the R4 and R5 region of the midgut.

To investigate further the role of *ninaD* in the whole *Drosophila* fly, as well as in the gut, we used *ninaD^1^* mutant. The *ninaD^1^* mutant allele is a base pair exchange (G to A), resulting in a stop codon at position 71 (Kiefer et al., 2002). In *Drosophila*, mutation in *ninaD* gene results in a carotenoid-free and consequently in vitamin A-deficient phenotype (Kiefer et al., 2002; Gu et al., 2004; Yang et al., 2007; Voolstra et al., 2006). *ninaD^1^* mutants has a reduced lifespan (Figure 1C) compared to wild-type CS, but longer than *jou*-deleted flies (*Del*). To determine whether the longevity effect in *ninaD^1^* mutants is prevented or enhanced by *jou*, we generated a *Del,ninaD^1^* double-mutant. Surprisingly, double-mutants has a longer lifespan than *jou*-deleted flies (*Del*) as well as *ninaD^1^*.

To confirm the RNAi knockdown efficacy, we measured by RT-qPCR the mRNA level of *ninaD* on dissected guts. Specific expression of UAS-ninaD-RNAi in enterocytes of *jou*-deleted flies is sufficient to restore (decrease to a roughly normal level) the mRNA level of *ninaD*, both with the Myo1A-Gal 4 driver (Figure 1D), as well as when it is expressed only in adulthood with the CG8997-GS line (Figure 1E). Moreover, RT-qPCR analysis showed that the mRNA levels of *ninaD* were hardly detectable in *ninaD^1^* mutants (Figure 1F). This decrease is probably caused by a rapid degradation of the mRNA of the *ninaD^1^* allele due to its nonsense mutation, as formerly hypothesized (Kiefer et al., 2002), as well as in a more general way for other mutated genes (Maquat, 2004). On the other hand, expression level of mRNA of *ninaD* in the double-homozygous mutants remains as high as (and even higher) in the *jou*-deleted flies (Figure 1F), suggesting that the putative degradation phenomena does not occur in *jou*-deleted flies. This result could suggest that the pseudouridylation of the ribosomal RNA by *jou*, as previously demonstrated (Soulé et al., 2020), or putatively of mRNA (McHahon et al., 2015) could prevent the degradation of the mutated *ninaD^1^* mRNA.

### *ninaD* is expressed in R4 and R5 segments of the adult midgut

To address the potential role and requirement of *ninaD*, the expression pattern of *ninaD* was determined more precisely in adult tissues. A high throughput analysis (FlyAtlas Anatomical Expression Data, *FlyBase*) has reported a weak mRNA expression of *ninaD* in various tissues in adult flies as eye, brain, and fat body, a low expression in the Malpighian tubules, head, and heart, and a moderate expression in the midgut. Nevertheless, its precise localization in this last tissue remain to be precisely determined. Unfortunately, since no antibodies against the *ninaD* protein exists, we carried out our histological analysis using the ninaD-GAL4 transgenic flies (Wang et al., 2007) to express the UAS-nls-GFP transgene. *ninaD* is faintly detected in some adult tissues, as the pericerebral fat body and retina (Figure 1G), while it is well detected in the midgut (Figure 1H). Higher magnification revealed that the expression in this tissue corresponds to two distinct areas, the R4 and R5 regions.

### *ninaD* is required in the midgut for sterol homeostasis

*ninaD* encodes a class B Scavenger receptor-like membrane protein homologous to those of mammalian, SR-BI and CD36 (Kiefer et al., 2002). In mammals, this receptor plays critical roles in cholesterol metabolism, mainly in the form of High-Density Lipoprotein cholesteryl ester (HDL-CE) (Acton et al., 1996). SR-BI also mediates the bidirectional flux of unesterified cholesterol between target cells and the circulating lipoproteins (HDL and LDL) (de la Llera-Moya et al., 1999; Endemann et al., 1993; Steinbrecher, 1999). In *Drosophila, ninaD* is essential for the cellular uptake of dietary carotenoids for the formation of the visual chromophore (Kiefer et al., 2002). However, a direct involvement of class B Scavenger receptors in insect cholesterol metabolism has not yet been reported. As *ninaD* is highly expressed in the R4 and R5 regions of the midgut and to determine whether *jou* mutants (in which *ninaD* is importantly upregulated) display defects in cholesterol homeostasis, we measured the cholesterol levels in 7 day-old adult controls and *jou*-deleted flies. *jou*-deleted flies have total cholesterol levels reduced by 14% (Figure 2A) with in particular a higher level of free cholesterol (Figure 2B), but a striking decrease in the amount of cholesterol-ester (Figure 2C). We assess whether the *ninaD^1^* mutation also interfere with cholesterol by quantifying these two metabolic parameters on *ninaD^1^* flies. The cholesterol-ester is importantly increased compared to control flies (Figure 2C), while the level of free cholesterol is decreased (Figure 2B), with a total cholesterol remaining unchanged (Figure 2A). Finally despite a high level of expression of a mutated form of *ninaD^1^* protein in the midgut, the overall level of the different forms of cholesterol were not different in *Del,ninaD^1^* double-mutants compared to *ninaD^1^* (Figure 2A, B, C). At first glance, these results seem to be in contradiction to a former reported study, in which they concluded that *ninaD^1^* adults have highly reduced carotenoid levels compared with wild-type, but have otherwise normal lipid contents (Kiefer et al., 2002), leading to suggest that *ninaD* was not involved in lipids metabolism. However, this previous study did not evaluate neither the level nor the different forms of cholesterol.

**Figure 2).**
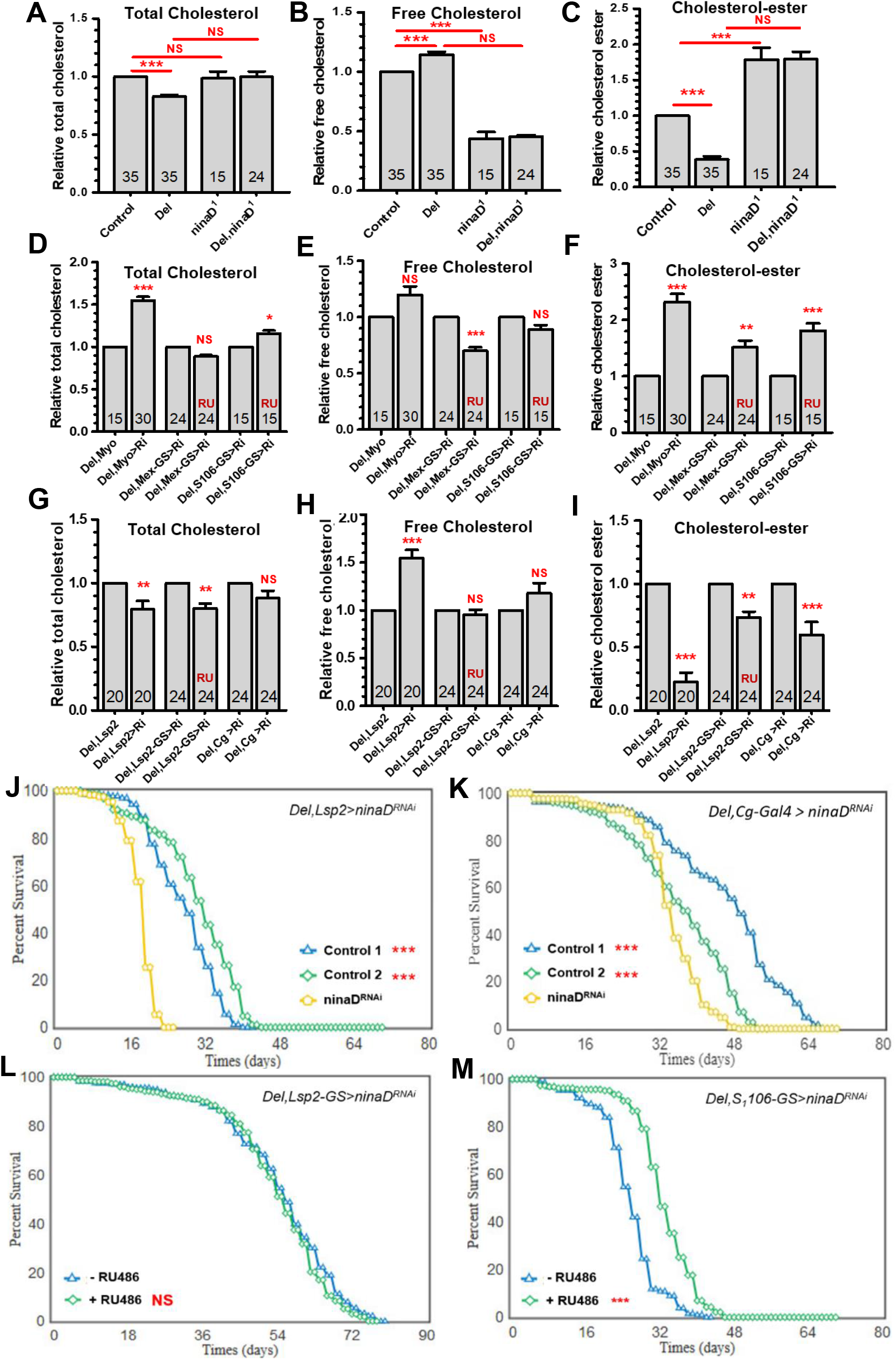
*ninaD* is involved in cholesterol homeostasis. **A-C)** Quantification of total **(A),** free **(B)**, cholesterol-ester **(C)** levels in wild-type controls (CS), *jou*-deleted (Del), *ninaD^1^*, and double-mutant (*Del,ninaD^1^*) flies. **D-F)** Quantification of various forms of cholesterol levels of flies expressing the UAS-ninaD-RNAi in the enterocytes of the *jou*-deleted flies by a Gal4 driver line (Del,Myo1A-Gal4), and by two conditional expression driver lines fed with RU486 only in adulthood (Del,Mex-GS and Del,S1106-GS). **G-H)** Quantification cholesterol levels of flies expressing the UAS-ninaD-RNAi in the fat body of the *jou*-deleted flies by two Gal4 driver lines (Del,Lsp2-GAL4 and Del,Cg-GAL4), and by a conditional expression driver line fed with RU486 only in adulthood (Del,Lsp2-GS). Numbers in the histograms = number of flies, errors bars = mean ± S.E.M (*p<0.05; **p<0.005; ***p<0.0005). p-value were calculated using the one-way ANOVA followed by a TUKEY test. **J-M)** Longevity test. Targeted expression of the UAS-ninaD-RNAi in the fat body in *jou*-deleted flies compared to their two respective controls (Control-1: Del,driver-Gal4/Del, Control-2 : Del,UAS-ninaD-RNAi), in two different driver-Gal4 lines: **J)** (Del,Lsp2-GAL4), **K)** (Del,Cg-Gal4). Conditional targeted expression of the UAS-ninaD-RNAi in the fat body in *jou*-deleted flies only in adulthood fed with RU486 compared to their controls (non-fed with RU486): **L)** (Del,Lsp2-GS), **(M)** (Del,S1106-GS) (***p<0.001). For detailed Statistics, see Table S2.

Since *ninaD* expression has been faintly detected in other tissues in adult flies, and especially in the fat body (Figure 1G and *Flybase*), we wonder if the expression of *ninaD* in this tissue is required and if its disruption could lead to cholesterol defects. We then perform a RNAi-mediated knockdown using various fat body and gut-specific drivers to express the UAS-ninaD-RNAi. In *jou-*deleted flies, restoring the *ninaD* mRNA level only in enterocytes using Myo1A-GAL4 restored the cholesterol-ester levels equivalent to those of wild-type. Whereas, the level of free cholesterol remains unchanged, while the total cholesterol is increased (Figure 2D, E, F). However, since *ninaD* is also importantly expressed during the development and especially in the larval midgut (*FlyBase*), we wonder if restoring its mRNA level in the epithelium of the gut, and so, only in adulthood is sufficient to restore the cholesterol defects observed in *jou-*deleted flies. We used the Mex-GS driver line (Soulé et al., 2020) to control the temporal expression of the UAS-ninaD-RNAi only in adulthood, by feeding flies with RU486, and thus quantify the levels of these compounds in adult. Again here, the various forms of cholesterol were rescued compared to control flies (Figure 2D, E, F). Similar results were obtained with the S1106-GS driver line, although this line is not expressed only in the epithelium of the midgut, but also in the fat body (Figure 2D, E, F). To conclude, in *jou*-deleted flies, in which the *ninaD* expression is importantly increased, the cholesterol defects can be rescued by the targeted expression of the ninaD-RNAi in enterocytes, allowing to rescue (decrease) the mRNA level of *ninaD*, indicating that these metabolic defects arises from a specific deregulation of *ninaD* function in this tissue.

### Downregulation of *ninaD*-mRNA in fat body in adulthood does not rescue cholesterol metabolism and viability

Since *ninaD* is also moderately expressed in adult head and weakly in adult fat body (Figure 1G and *FlyBase*), we wonder if *ninaD* functions in this tissue is required for maintaining proper metabolic homeostasis. To test this hypothesis, we rescue in *jou-*deleted flies, the *ninaD* mRNA level by expressing the UAS-ninaD-RNAi in the fat body. Cholesterol levels were not restored in *jou-*deleted flies upon expression of ninaD-RNAi exclusively in the fat body with Lsp2-GAL4 or Cg-GAL4 drivers (Figure 2G, H, I). Since the Lsp2-GAL4 line is also expressed at different developmental stages, we use a conditional adult-specific expression line, the Lsp2-GS (Ragheb et al., 2017), to restrict the expression of the ninaD-RNAi into the fat body only in adulthood. As for the Lsp2-GAL4, the expression of the ninaD-RNAi did not rescue the *jou* phenotypes, since the different forms of cholesterol were not restored compared to the non-induced flies (Figure 2G, H, I).

In complement, we also assessed the adult lifespan of these different fat body Gal4 lines. Unexpectedly, in *jou-*deleted flies, targeted knockdown of mRNA level of *ninaD* in the fat body shortens lifespan, both for Del,Lsp2-Gal4 (Figure 2J), and Del,Cg-Gal4 (Figure 2K). However, the expression of ninaD-RNAi only in adulthood, by the Lsp2-GS line did not affect the lifespan (Figure 2L), suggesting that the deleterious effect of the ninaD-RNAi might result from its expression throughout developmental stages. Finally, the targeted expression of the ninaD-RNAi both into the epithelium of the gut and the fat body, and so, only in adulthood (Del,S1106-GS) increases lifespan (Figure 2M), indicating that the rescue of a homeostatic *ninaD* mRNA level is required in the gut. In brief, these results indicate that the essential functions of *ninaD* reside in the midgut for maintaining the cholesterol homeostasis.

### In the gut, loss of *ninaD* decreases the free cholesterol

The homology of *ninaD* to SR-BI suggests that these proteins may have conserved biochemical function (de la Llera-Moya et al., 2001; Connelly et al., 2001; Herboso et al., 2011). Nonetheless, many important questions concerning the precise function of *ninaD* remain unanswered. Some studies have pointed out the role of various genes in the cholesterol homeostasis in the gut, as *NPC1 & 2* (Huang et al., 2007; Voght et al., 2007), *magro*, a secreted Lipase A (Sieber et al., 2012), or DHR96, a nuclear receptor that regulates the cholesterol homeostasis (Horner et al., 2009). However, it remains unclear whether *ninaD* specifically promotes the absorption of sterols or whether this protein participates in the intracellular trafficking of cholesterol in the intestinal epithelium. As a first step, we determine, at the cellular level, the precise localization of *ninaD* in enterocytes. Since unfortunately, no antibody exists against *ninaD*, we target a *ninaD* transgene coupled to EGFP (UAS-ninaD-EGFP) (Gomez-Diaz et al., 2016) to the midgut using two independent intestinal lines, Mex-Gal4 and Myo1A-Gal4. The similar results obtained with both lines revealed that the ninaD-EGFP fusion protein is specifically localized in the plasma membrane of enterocytes (Figure 3A, B). Based on our observations that upregulation of *ninaD* in *jou*-deleted flies affects the sterol metabolism, we next examined the sterols distribution in the midgut using filipin staining (Huang et al., 2007; Voght et al., 2007). Filipin is a fluorescent polyene macrolide that binds 3-β-hydroxysterols such as free cholesterol. Unesterified (free) cholesterol in the plasma membrane of enterocytes is decreased in *jou*-deleted flies relative to wild-type controls (Figure 3C, D). Similarly, filipin stainings are also decreased in *ninaD^1^* midgut, as well as in double-mutant (*Del,ninaD^1^*). A quantification of fluorescence confirms these results (Figure 3G).

**Figure 3).**
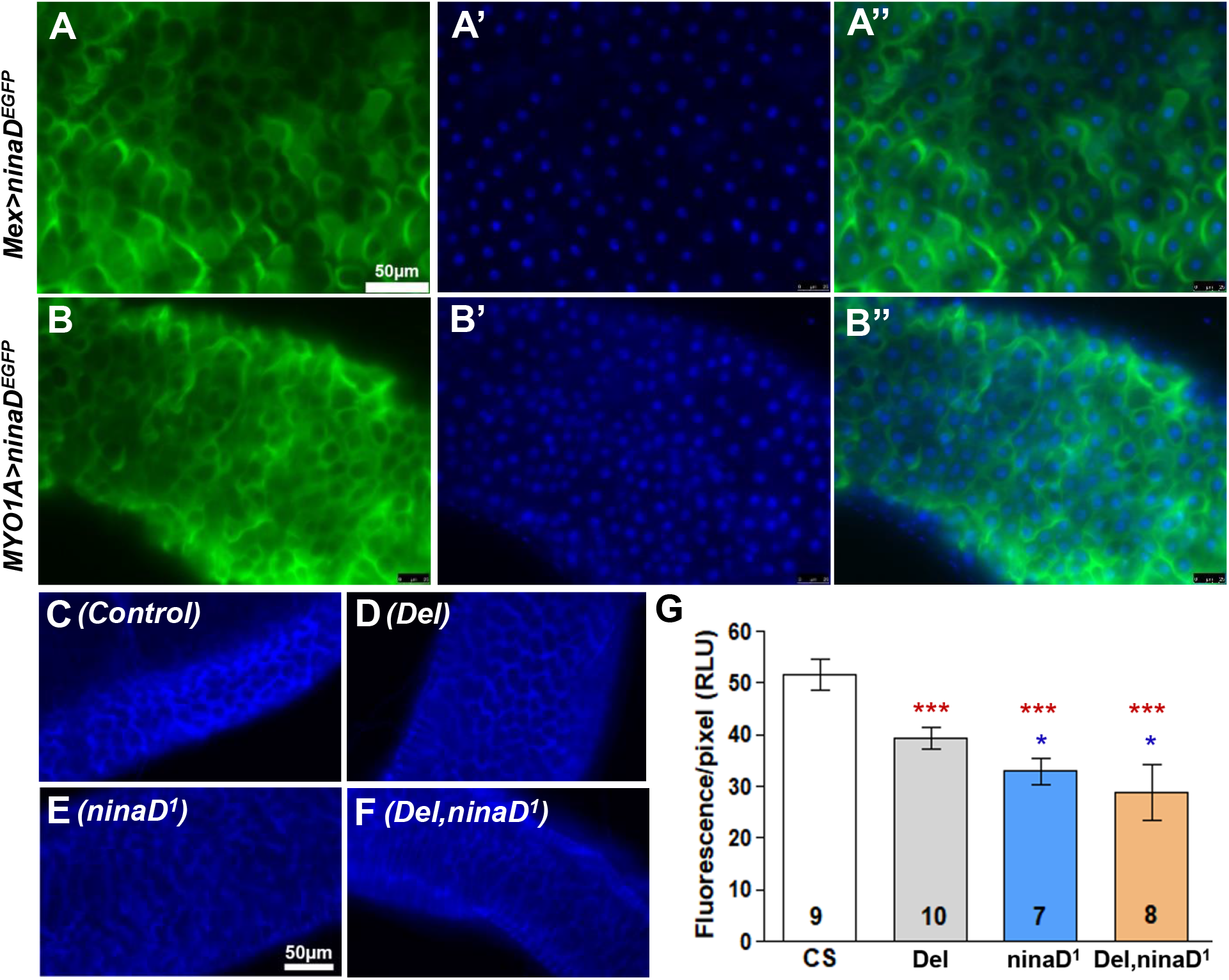
*ninaD* is localized in plasma membrane of enterocytes. **A)** Mex-GAL4>UAS-ninaD-EGFP. A: GFP (green), A’: DAPI staining (blue) and A’’: overlay. **B)** MYO1A-GAL4>UAS-ninaD-EGFP. B: GFP (green), B’: DAPI staining (blue) and B’’: overlay. **C-F)** Filipin staining revealing the unesterified (free) cholesterol in plasma membrane of enterocytes. **C)** Control (CS), **D)** *jouvence*-deleted (*Del*), **E)** *ninaD^1^*, **F)** double-mutant (*Del,ninaD^1^*). **G)** Fluorescence quantification of filipin of the gut for the four genotypes.

### Deregulation of *ninaD* in the gut leads to neurodegenerative lesions in old flies

In *Drosophila*, impairment in cholesterol homeostasis has already been associated with various neurodegenerative disorders, as in Niemann-Pick disease (Huang et al., 2007; Voght et al., 2007; Philips et al., 2008), as well as in *löchrig* mutant encoding a gene which interferes with cholesterol homeostasis (Tschäpe et al., 2002). In human, altered cholesterol metabolism and hypercholesterolemia significantly contribute to neuronal damage and progression of Alzheimer disease (Gamba et al., 2015; Fernandez-Pérez et al., 2018). These previous reports suggest that the nervous system is highly sensitive to alterations in cholesterol homeostasis. This sensitivity is tough to result from the high metabolic demand of neurons and their supporting cells. Here, since neurodegeneration is predicted to be age progressive, we examined the brain of our different studied genotypes in order to assess, in old flies (40-days), neurodegeneration as manifested by the appearance of vacuolar lesions. At 5-days-old, wild-type controls display an intact central brain with no detectable vacuole (Suppl. Figure 1). However, at 40-days-old, wild-type flies show clear signs of neurodegeneration, with the appearance of several vacuoles of different sizes (Figure 4A). In 40-days-old *jou*-deleted flies, vacuolization is more severe in the central brain (Figure 4B), which is more pronounced in number and affected size-area than age-matched wild-type controls (Figure 4K, L). The quantification of the number of vacuoles (Figure 4K), as well as the surface area of the lesions (Figure 4L), is significantly higher in *jou*-deleted flies than in controls. Similarly, at 40-days-old, *ninaD^1^* and *Del*, ninaD^1^ double-mutant show an increase in vacuolization relative to wild-type controls (Figure 4C, D, K, L).

**Figure 4).**
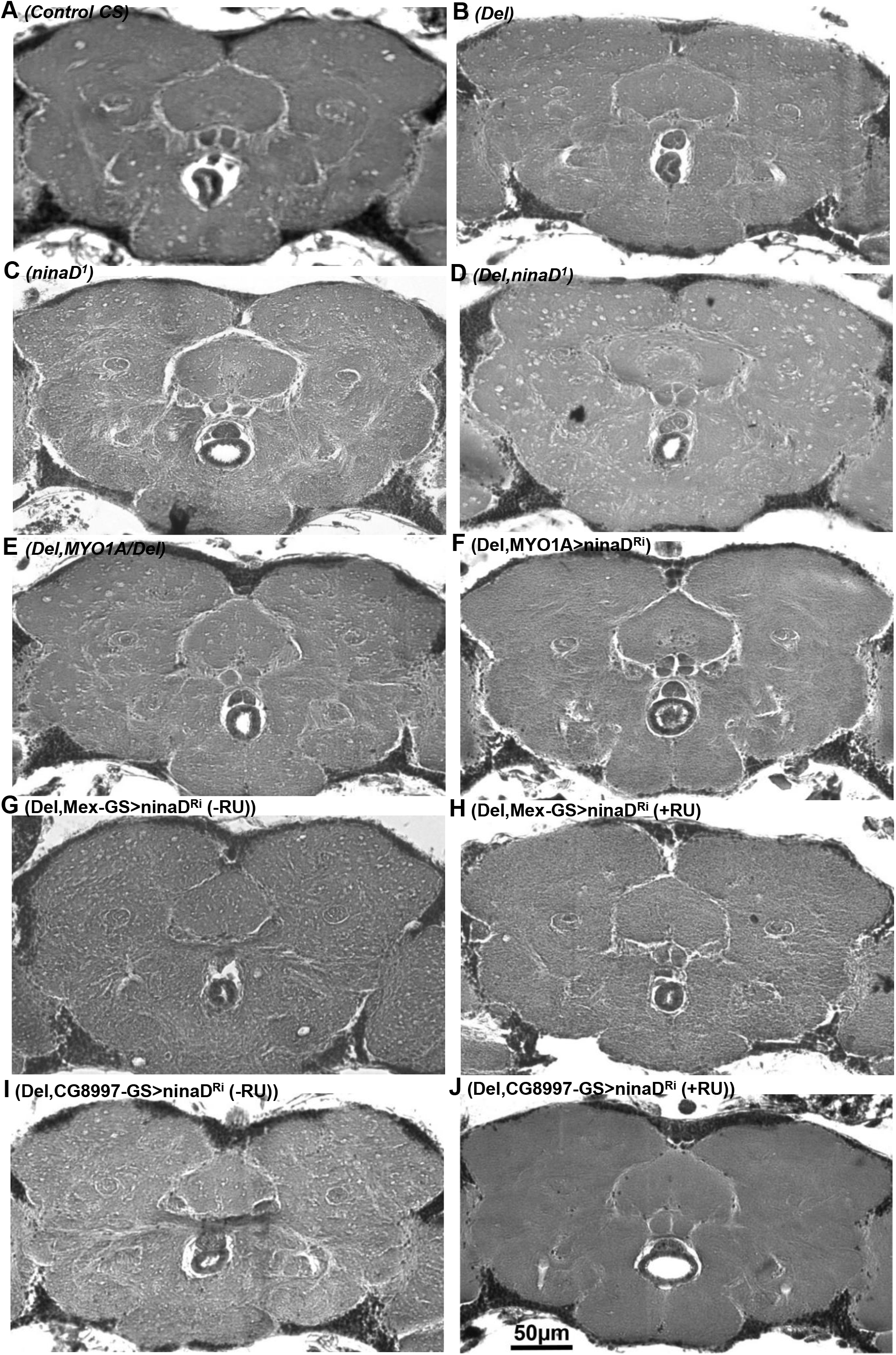
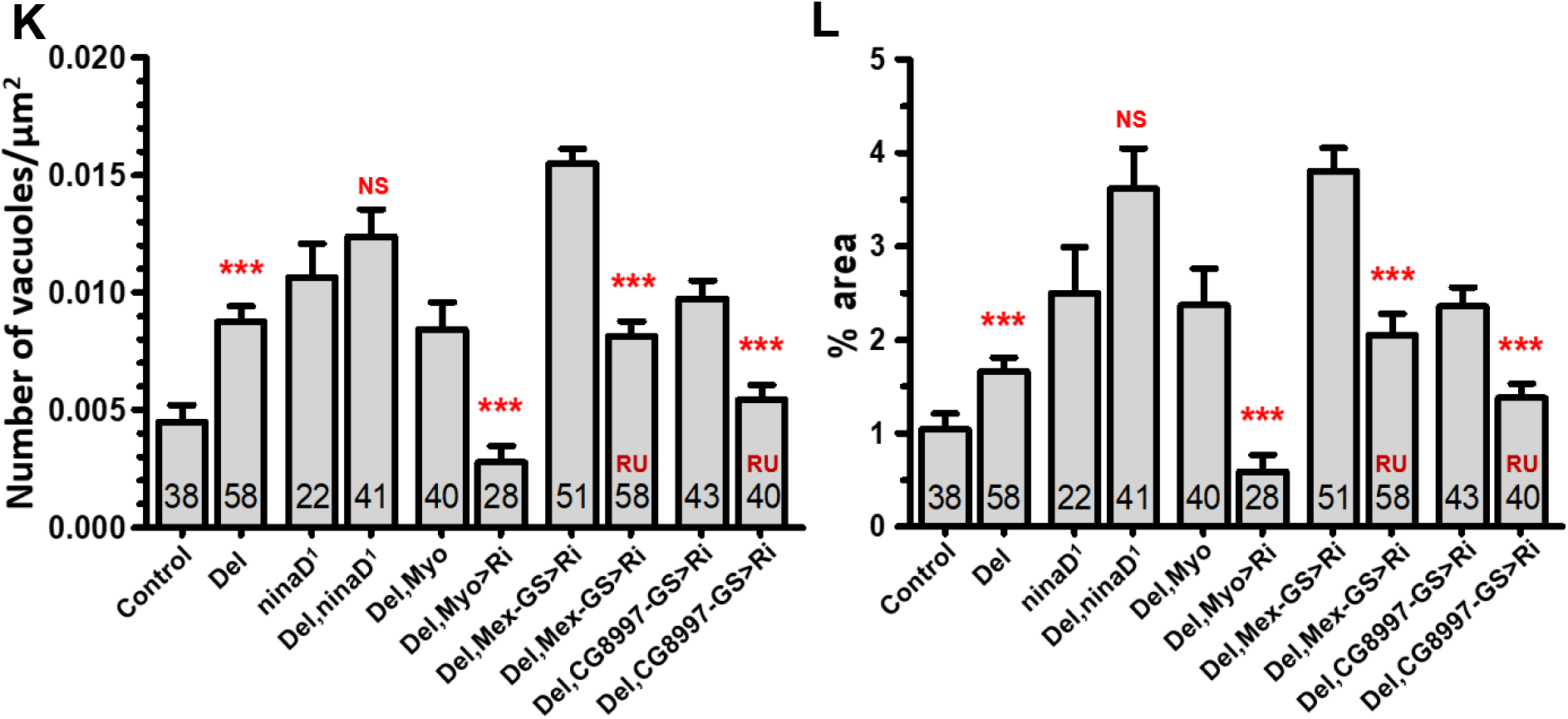
Rescuing ninaD-mRNA level in the gut protects against neurodegenerative lesions in old flies. **A-J)** 7 μm paraffin sections of the brain revealing the neurodegenerative lesions (vacuoles) in the brain. **A)** Control (CS), **B)** *jou*-deleted (Del), **C)** *ninaD^1^*, **D)** *Del,ninaD^1^*, **E)** Del,MYO1A-GAL4/Del (control), **F)** Del,MYO1A-GAL4>UAS-ninaD-RNAi, **G-H)** Del,Mex-GS>UAS-ninaD-RNAi (No RU) (G), and (+RU) (H), **I-J)** Del,CG8997-GS>UAS-ninaD-RNAi (No RU) **(I)**, and (+RU) **(H)** (scale bar=50 μm for all images). **K)** Number of vacuoles for genotypes analyzed above. **L)** Area of vacuoles (in %) compared to the analyzed surface area of the brain. Numbers in histograms = number of flies, errors bars = mean ± S.E.M, *p<0.05; **p<0.005; ***p<0.0005). p-value were calculated using the one-way ANOVA followed by a TUKEY test.

Since we have demonstrated that *ninaD* is required for sterol metabolism from the midgut epithelium, we wonder if the downregulation of *ninaD* in the midgut (the rescue) might rescue the age-dependent brain deterioration by restoring cholesterol levels. In *jou*-deleted flies expressing the ninaD-RNAi driven by Myo1A-GAL4 in the midgut, the neurodegenerative lesions are rescued (Figure E, F), as well as the number of vacuoles and the affected area of the brain (Figure 4K, L). Even more precisely, the targeted expression of the ninaD-RNAi in the midgut, but restricted only in adulthood, and so by two independent lines, Mex-GS (Figure 4G, H) and CG8997-GS (Figure 4I, J), rescue the neurodegenerative lesions in old flies (Figure 4K, L). Altogether, these results indicate that restoring the mRNA level of *ninaD* in the midgut was sufficient to mitigate the mutant effects, and serves as a neuroprotective role.

Recently, we have determined the role of *jouvence* in human, using culture cells. The overexpression of *jou* increases the proliferation of the cells, whereas, in contrast, its knockdown, by siRNA, decreases it (El-Khoury et al., 2020). To assess its molecular mechanisms, transcriptomic analysis have been performed on up- and down-regulation of *jou*, revealing that about 6000 genes are deregulated in each condition. The KEGG analysis indicated that the metabolic pathways are the main deregulated pathways. An advanced analysis revealed that among them, the HMG-CR (3-hydroxy-3-methylglutaryl-CoA reductase), the limiting enzymes involved in cholesterol synthesis (which is also the target of statins, the human anti-cholesterol medications), follows the up- and down-expression level of *jou*, suggesting that this gene is likely directly controlled by *jou*. Interestingly, some snoRNAs have already been involved in stress responses and metabolic homeostasis. The snoRNAs U32a, U33, and U35a are critical mediators of oxidative stress (Michel et al., 2011), while the snoRNA U60 regulates intracellular cholesterol trafficking between the plasma membrane and the endoplasmic reticulum (Brandis et al., 2013). Similarly, the snoRNA U17 regulates cellular cholesterol trafficking through the hypoxia-upregulated mitochondrial regulator (HUMMR) (Jinn et al., 2015). Therefore, we hypothesize that one of the main cellular effect of *jou* in human cells could be due to the regulation of cholesterol homeostasis. Thus, it is compelling to observe that *jou* seems to lead to similar cellular effects in *Drosophila* and human.

## LIMITATIONS OF STUDY

Since both in *ninaD* mutant as well as in *jou*-deleted flies we observe a perturbation of cholesterol-ester but not of total cholesterol, it suggests that a *ninaD* regulates the balance between free and cholesterol-ester. Thus, *ninaD* seems to be rather involved in the de-esterification process of cholesterol, instead of in its absorption. Future experiments will be required to determine precisely the molecular mechanism by which *ninaD* regulates the different forms of cholesterol. Moreover, although restoring the level of cholesterol-ester is sufficient to rescue the neurodegenerative lesions in old flies, more investigations will be required to the exact and precise nature of the neurodegenerative (vacuoles) lesions.

## MATERIALS AND METHODS

### *Drosophila* lines

*Drosophila melanogaster* flies were grown on standard medium (1.04% agar, 8% cornmeal, 8% brewer yeast and 2% nipagin as a mold inhibitor) at 25°C, 12:12 light:dark cycle in a humidity-controlled incubator. For ageing experiments, to avoid larval growth and overpopulation in vials, 15 adult flies per vial were transferred to new food vials every 2 days. *Canton-S* (*CS*) flies were used as control. The results described in this present study were obtained from females. All genotypes were outcrossed to the wild-type CS (cantonization) at least 6 times to thoroughly homogenize the genetic background. RU486 (Mifepristone) (*Sigma-Aldrich*, Cat. #M8046) to induce the Gene-Switch activity was dissolved in ethanol and mixed into the media when preparing food vials. RU486 doses used were 25 μg.mL^−1^ final concentration. *S1106-GS*, and *P{ninaD-GAL4.W}3* were obtained from the Bloomington *Drosophila* Stock Center (http://flystocks.bio.indiana.edu/). *UAS-ninaD-RNAi* line (NIG.31783R) was obtained from NIG-Fly (Japan, https://shigen.nig.ac.jp/fly/nigfly/). Myo1A-Gal4 was kindly provided by B.A. Edgar (Heidelberg, Germany), Mex-Gal4 from G.H. Thomas (Pennsylvania, USA), *CG8997-GS* from H. Tricoire (Paris, France), *ninaD^1^* mutant from D. Vasiliauskas (Gif-sur-Yvette, France), *P{UAS-ninaD-EGFP}* from R. Benton (Lausanne, Switzerland), and *UAS-nls-GFP*(*4776*) from F. Rouyer (Gif-sur-Yvette, France). These transgenic lines were then introduced into the *jouvence* deletion background (*Del*) by standard genetic crosses. Mex-GS driver line have been generated in our laboratory (Soulé et al., 2020). *Del,ninaD^1^* double-mutant has been generated by recombination since they are located both on the second chromosome. In brief, a cross between the *jouvence*-deleted flies and the *ninaD^1^* null mutant has been performed. Subsequent progeny were analyzed by PCR on the genomic DNA, followed by sequencing the PCR products to search for the presence of the nonsense mutation of *ninaD^1^* mutant (genotyping).

### Lifespan analysis

Following amplification, flies were harvested every day after hatching. Female flies were maintained with males in fresh food vials for 4 days at a density of 30 individuals per vial. On day 4, males were removed, while females were placed in a cage (about 200 to 300 females per cage), as in Soulé et al. (2020). Cages (100 mm high and 60 mm diameter = 283 cm^3^) used for lifespan experiments are made in plexiglass. The top is covered by a mesh allowing a good aeration, while the bottom (floor) is closed by a petri dish containing the food medium. The aging animals were transferred to fresh food every two days, by changing the petri dish, and the number of dead flies was scored. The lifespan plots were generated by calculating the percentage of survivorship every two days and plotting viability as a function of time (days) using log-rank test (Yang et al., 2011).

### Quantitative RT-PCR

RNAi knock-down efficiency was measured on dissected midguts or whole flies from 7-days-old female flies. 30-40 midguts per sample were immediately frozen in nitrogen liquid and kept at −80°C. Each genotype was collected in triplicate. Total RNA from collected midguts were extracted using SV Total RNA Isolation System kit (*Promega*, Cat. #Z3100) according to the manufacturer’s protocol. RNA concentration was measured using a spectrophotometer (*BioRad*) and purity of the samples was estimated by OD ratios (A_260_/A_280_ ranging within 1.8-2.2). Total RNA were then analyzed on 1.5% agarose gel stained with ethidium bromide to verify any DNA contamination. All the samples were processed for 35 PCR cycles (95°C for 45 s; 55 - 60°C for 30 s; 72°C for 30 s), with a final extension step at 72°C for 10 min. cDNA were transcribed from 1 μg DNA-free total RNA using the M-MLV reversible transcriptase (*Invitrogen*, Cat. #28025-013) and oligo(dT) primer (*Promega*, Cat. #C1101). Quantitative RT-PCR was performed on a Bio-Rad CFX96 instrument (*BioRad*) using LightCycler^®^ FastStart DNA Master^PLUS^ SYBR Green I (*Roche*, Cat. #03752186001). All assays were done in triplicate. Data were analyzed according to the ΔΔCt method, and normalized to *α-Tub84B* levels. Primers used are summarized in the Supplemental Experimental Procedures (see Supplementary Table S1).

### Brain histology

Flies were filled in collars and fixed for 4h in fresh Carnoy’s fixative (ethanol:chloroform:acetic acid at 30:15:5) (Belgacem et al., 2006). Heads were then embedded into paraffin for sectioning using standard histological procedures. Sections of 7 μm were stained with 1% Toluidine blue (*Sigma-Aldrich*, Cat. #89640-5G), and 1% Borax for 8 min. Samples were mounted with Mowiol, and photographed on a Leica DM600B light microscope (*Leica*, Germany) equipped with a Hamamatsu C10600 ORCA-R^2^ digital camera. Quantification of the neurodegenerative lesions was performed on half of the brains using ImageJ software. Briefly, images were converted to 32 bits and a minimum threshold have applied to the entire image. The area above the threshold, whose correspond to the vacuoles (neurodegenerative lesions), were then labeled and quantified in the brain to estimate their number and size. Data are expressed as number of vacuolar lesions per μm^2^ of measured surface.

### Filipin staining

Midguts from female flies at 7 day-old were fixed in 4% PFA for 30 min, and rinsed twice in 1X PBS + 0.05% Tween 20 (PBST). A fresh 2 mg.mL^−1^ stock solution of filipin (*Sigma-Aldrich*, Cat. #F4767-1MG) in 100% ethanol was used. Samples were stained in the dark with 50 μg.mL^−1^ filipin diluted in PBST solution for 30 min, washed with PBST, and mounted in Mowiol. Stained tissues were immediately visualized with a Leica DM 600B light microscope (Leica, Germany), and imaged with a Hamamatsu C10600 ORCA-R^2^ digital camera, using an excitatory wavelength of 360 - 480 nm.

### Cholesterol measurement

Metabolic cholesterol quantification, measured as described in (Li et al., 2017) with some slight modifications, were carried out on 24 single 7-days-old adult female flies. Flies were weighted and rinsed twice with PBS to remove all remaining food and dirtiness on their cuticle. In brief, animals were homogenized in a Potter apparatus with a 7:11:0.1 chloroform/isopropanol/NONIDET P40 solution using a hand homogenizer. After a 10 min spin at 13,000g the supernatant was dried in a Speed Vacuum Concentrator for 2h at 50°C. The lyophilized material was then dissolved in 1X reaction buffer and split into two samples. 0.5 μL cholesterol esterase was added to one sample and both samples were incubated at 37°C overnight. Free cholesterol and cholesterol ester levels were measured using the Amplex Red cholesterol assay kit (*Invitrogen*, Cat. #A12216), according to the manufacturer’s instructions. Fluorescence was read using a Tecan™ GENios^®^ plate reader (*TECAN*) with a 530/590 nm filter set and compared to a standardization curve.

### Analysis of expression pattern of *ninaD*

All histological experiments were carried out on dissected guts and cryosections of whole flies from adult females aged of 7 days. To determine the expression pattern of *ninaD* (ninaD-GAL4>UAS-nls-GFP, and Mex-GAL4>UAS-ninaD-EGFP), samples were fixed in 4% PFA for 30 min, washed three times with PBS, and mounted in Mowiol. For dissected guts, tissues were counterstained with DAPI (*Roche*, Cat. #10236276001) for 5 min in the dark. Images were collected using a Leica DM 600B light microscope (*Leica*, Germany), equipped with a Hamamatsu C10600 ORCA-R^2^ digital camera.

### Statistical analyses

Statistical comparisons were done with GraphPad Prism. Data were analyzed using one-way ANOVA test. Significance levels in figures were represented as *p < 0.05, **p < 0.01, ***p < 0.001. All quantitative data are reported as the mean ± S.E.M. (Standard Error of the Mean). Lifespan assays were subjected to survival analysis (log-rank test) using the freely available OASIS software (Yang et al., 2011).

## ACKNOWLEDGMENTS

We thank L. Mellottée for her technical assistance. J. Montagne (I2BC, CNRS, Gif-sur-Yvette) for the critical reading of the manuscript and for helpful discussion. H. Tricoire (Paris, France), B.A. Edgar (Heidelberg, Germany), G.H. Thomas (Pennsylvania, USA), D. Vasiliauskas (Gif-sur-Yvette, France), R. Benton (Lausanne, Switzerland), F. Rouyer (Gif-sur-Yvette) and J.E. O’Tousa (Indiana, UAS) for fly stocks. This work was supported by IRS IDEX Brainscopes, and by the CNRS (France).

## AUTHOR CONTRIBUTIONS

JRM and SS conceived and designed the project. SS and JRM performed and analyzed the experiments. SS and JRM discussed the results and wrote the manuscript.

## DECLARATION OF INTERESTS

The authors declare no competing interests.

## Notes

### Competing Interest Statement

The authors have declared no competing interest.

