## Supplementary Information (Tables and Figure) for "*ninaD* regulates cholesterol homeostasis from the midgut which protects against neurodegeneration"

### Table S1) RT-qPCR primers used.

| Gene Name | Primer Forward (5' - 3') | Primer Reverse (5' - 3') |
|---|---|---|
| *α-Tub84B* | TGTCGCGTGTGAAACACTTC | AGCAGGCGTTTCCAATCTG |
| *ninaD* | ACCAAATGCGGAATAGCAAC | GGCGTAATGCAAAAATTCGT |

### Table S2) Statistics of longevity tests.

| **Figure 1A** | **Age in days at % mortality** | | | | | | |
|---|---|---|---|---|---|---|---|
| **Genotype** | **Number of flies** | **25%** | **50%** | **75%** | **90%** | **100%** | **P-value** |
| Del,MYO1A-GAL4 $^{(1)}$ (Control 1) | 312 | 26 | 28 | 32 | 36 | 40 | $0^{(1)}$ $0^{(2)}$ |
| Del,UAS-ninaD$^{RNAi\ (2)}$ (Control 2) | 275 | 26 | 32 | 38 | 40 | 44 | |
| Del,MYO1A-GAL4>UAS-ninaD$^{RNAi}$ (ninaD$^{RNAi}$) | 231 | 44 | 48 | 50 | 52 | 54 | |
| **Figure 1B** | | | | | | | |
| Del,CG8997-GS>UAS-ninaD$^{RNAi}$ (- RU486) | 191 | 29 | 35 | 41 | 45 | 63 | 0 |
| Del,CG8997-GS>UAS-ninaD$^{RNAi}$ (+ RU486) | 208 | 50 | 56 | 58 | 62 | 70 | |
| **Figure 1C** | | | | | | | |
| Control (CS) | 216 | 29 | 31 | 33 | 35 | 37 | $0^{(1)}$ $0^{(2)}$ $0^{(3)}$ |
| Del $^{(1)}$ | 254 | 17 | 21 | 23 | 25 | 27 | |
| ninaD$^{1\ (1)\ (2)}$ | 195 | 23 | 29 | 31 | 35 | 37 | |
| Del,ninaD$^{1\ (1)\ (2)\ (3)}$ | 227 | 39 | 47 | 55 | 61 | 71 | |
| **Figure 2M** | | | | | | | |
| Del,Lsp2-GAL4 $^{(1)}$ (Control 1) | 258 | 22 | 28 | 34 | 36 | 40 | $0^{(1)}$ $0^{(2)}$ |
| Del,UAS-ninaD$^{RNA\ (2)i}$ (Control 2) | 275 | 26 | 32 | 38 | 40 | 44 | |
| Del,Lsp2-GAL4>UAS-ninaD$^{RNAi}$ (ninaD$^{RNAi}$) | 276 | 17 | 19 | 21 | 21 | 23 | |
| **Figure 2N** | | | | | | | |
| Del,Cg-GAL4 $^{(1)}$ (Control 1) | 236 | 37 | 49 | 55 | 63 | 67 | $0^{(1)}$ $0^{(2)}$ |
| Del,UAS-ninaD$^{RNAi\ (2)}$ (Control 2) | 300 | 29 | 39 | 47 | 49 | 53 | |
| Del,Cg-GAL4>UAS-ninaD$^{RNAi}$ (ninaD$^{RNAi}$) | 296 | 31 | 35 | 39 | 43 | 49 | |
| **Figure 2O** | | | | | | | |
| Del,Lsp2-GS>UAS-ninaD$^{RNAi}$ (- RU486) | 277 | 46 | 56 | 64 | 70 | 78 | 0.0583 |
| Del,Lsp2-GS>UAS-ninaD$^{RNAi}$ (+ RU486) | 278 | 48 | 56 | 62 | 68 | 76 | |
| **Figure 2P** | | | | | | | |
| Del,$S_1$106-GS>UAS-ninaD$^{RNAi}$ (- RU486) | 212 | 22 | 26 | 28 | 34 | 42 | 0 |
| Del, $S_1$106-GS>UAS-ninaD$^{RNAi}$ (+ RU486) | 187 | 30 | 32 | 38 | 40 | 46 | |

**Suppl. Figure 1)**

A 7 µm paraffin section of a young (5 day-old) brain wild-type fly. The brain is uniform, and without vacuoles.

Suppl. Figure 1 Soulé & Martin

**Control (CS) - Young**

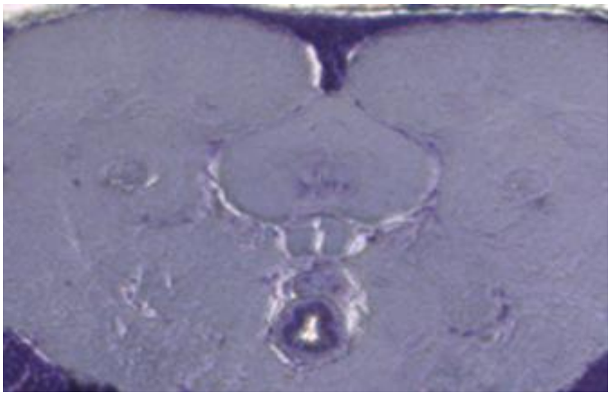
